# Quantification of gallium cryo-FIB milling damage in biological lamella

**DOI:** 10.1101/2023.02.01.526705

**Authors:** Bronwyn A. Lucas, Nikolaus Grigorieff

## Abstract

Cryogenic electron microscopy (cryo-EM) has the potential to reveal the molecular details of biological processes in their native, cellular environment at atomic resolution. However, few cells are sufficiently thin to permit imaging with cryo-EM. Thinning of frozen cells to <500 nm lamellae by cryogenic focused ion beam (FIB) milling has enabled visualization of cellular structures with cryo-EM. FIB-milling represents a significant advance over prior approaches because of its ease of use, scalability, and lack of large-scale sample distortions. However, the amount of damage caused by FIB-milling to the generated thin cell section has not yet been determined. We recently described a new approach for detecting and identifying single molecules in cryo-EM images of cells using 2D template matching (2DTM). 2DTM is sensitive to small differences between a molecular model (template) and the detected structure (target). Here we use 2DTM to demonstrate that under the standard conditions used for machining lamellae of biological samples, FIB-milling introduces a layer of variable damage that extends to a depth of 60 nm from each lamella surface. This thickness exceeds previous estimates and limits the recovery of information for *in situ* structural biology. We find that the mechanism of FIB-milling damage is distinct from radiation damage during cryo-EM imaging. By accounting for both electron scattering and FIB-milling damage, we find that FIB-milling damage will negate the potential improvements from lamella thinning beyond 90 nm.

**Significance:** The molecular mechanisms of biological macromolecules and their assemblies is often studied using purified material. However, the composition, conformation and function of most macromolecules depend on their cellular context, and therefore, must also be studied inside cells. Focused ion beam (FIB) milling enables cryogenic electron microscopy to visualize macromolecules in cells at close to atomic resolution by generating thin sections of frozen cells. However, the extent of FIB-milling damage to frozen cells is unknown. Here we show that Ga^+^ FIB-milling introduces damage to a depth of ∼60 nm from each lamella surface, leading to a loss of recoverable information of up to 20% in 100 nm samples. FIB-milling with Ga^+^ therefore presents both an opportunity and an obstacle for structural cell biology.

## Introduction

Cryogenic electron microscopy (cryo-EM) has enabled visualization of purified macromolecular complexes at atomic resolution (1, 2). A more complete understanding of molecular function requires visualizing their location, structure, and interactions in the native cellular environment. The internal architecture of cells can be preserved with high fidelity by vitrification allowing for the visualization of molecules at high resolution directly in the cell (*in situ*) with cryo-EM (1). However, with few exceptions, cells are too thick to be electron-transparent, and therefore require thinning.

Cryo-EM of vitreous sections (CEMOVIS) was an initial solution to generating thin slices of high-pressure frozen cells using a cryo-ultramicrotome (2). However, the process requires a skilled user, is difficult to automate and introduces compression artifacts, which together have limited the widespread utility of this approach (3).

Focused ion beam (FIB) milling is a technique in common use in materials science that has been adapted to produce thin cell sections for *in situ* cryo-EM under cryogenic conditions (4–6). In place of a physical ultramicrotome, a focused beam of ions, typically produced from a gallium liquid metal ion source (LMIS), is used to sputter material above and below a thin section of the cell known as a lamella (6). FIB-milling has higher throughput relative to CEMOVIS because of its ease of use, commercial availability, and computational control allowing for automation of lamella production (7, 8). As a result, cryo-FIB milling for TEM lamella preparation of cells has recently seen widespread adoption and is now the predominant method for preparing cells for *in situ* cryo-EM (9).

It has been demonstrated recently that it is possible to generate near-atomic resolution reconstructions by averaging subtomograms from vitreously frozen cells (10, 11). These successes highlight the need for a more quantitative understanding of potential sample damage introduced during FIB-milling that could limit both the resolution of *in situ* reconstructions and the ability to accurately localize molecules in cells.

Organic materials are particularly sensitive to radiation damage upon interaction with high-energy particles. Simulations of the stopping range in matter (SRIM) of ions in a glancing incidence beam at 30 keV, the typical conditions for cryo-lamella preparation for TEM, will implant Ga+ ions in frozen cells to a depth of 20-30 nm (5, 12). After accounting for removal of ∼10 nm of material due to the concurrent milling action, the implantation zone is anticipated to be ∼10-20 nm from the lamella surface (5). Cascading atomic collisions between Ga^+^ ions and sample atoms as the Ga^+^ ions imbed in the sample will introduce additional damage to an unknown depth from each lamella surface (13). Such damage introduced during FIB-milling would decrease the usable volume of a lamella and could limit the resolution of *in situ* determined structures. Cryo-electron tomography (cryo-ET), the predominate method for characterizing molecular structure *in situ*, generates tomograms that are limited in resolution to about 20 Å (14), and is therefore not suitable for measuring differences in individual particle quality at high-resolution.

We have recently described a new approach, 2D template matching (2DTM) (15), to locate molecular assemblies in three dimensions with high precision in 2D cryo-EM images of unmilled cells (16, 17) and FIB-milled lamellae (18). Cross-correlation of a high-resolution template generated from a molecular model with a cryo-EM image produces a 2DTM signal-to-noise ratio (SNR) that reflects the similarity between the template and the individual target molecules in the image (15–18).

In the present study, we apply 2DTM to quantify target integrity within FIB-milled lamellae at single molecule resolution. We find that FIB-milling appreciably reduces target integrity to a depth of ∼60 nm from the lamella surface, more than previously appreciated. We find that the nature of FIB-milling damage is distinct from electron radiation damage, consistent with inter-atomic collisions, rather than electronic interactions, being responsible for the damage. By comparing signal loss due to FIB-milling damage to signal loss in thick samples due to inelastic electron scattering and molecular overlap, we show that recovery of structural information in 100 nm lamellae is reduced by ∼20%.

## Results

### FIB-milling introduces a layer of reduced structural integrity

A 2DTM template represents an ideal, undamaged model of the molecule to be detected. Any damage introduced during FIB-milling will therefore decrease the correlation with the undamaged template, leading to a lower 2DTM SNR. Ribosomes are present at high density and relatively evenly distributed in the cytoplasm of the yeast *Sacchromyces cerevisiae* (18), and therefore present an ideal 2DTM target to quantify differences in target integrity. We prepared FIB-milled lamellae of *S. cerevisiae* cells of thickness varying from 120 nm to 260 nm (**Fig. 1*A* and *B***). In 30 images of the yeast cytoplasm from four lamellae, we located 11030 large ribosomal subunits (LSUs) using 2DTM (**Fig. 1*C* and *D, SI Appendix*, Fig. S1*A* and *B***).

**Figure 1:**
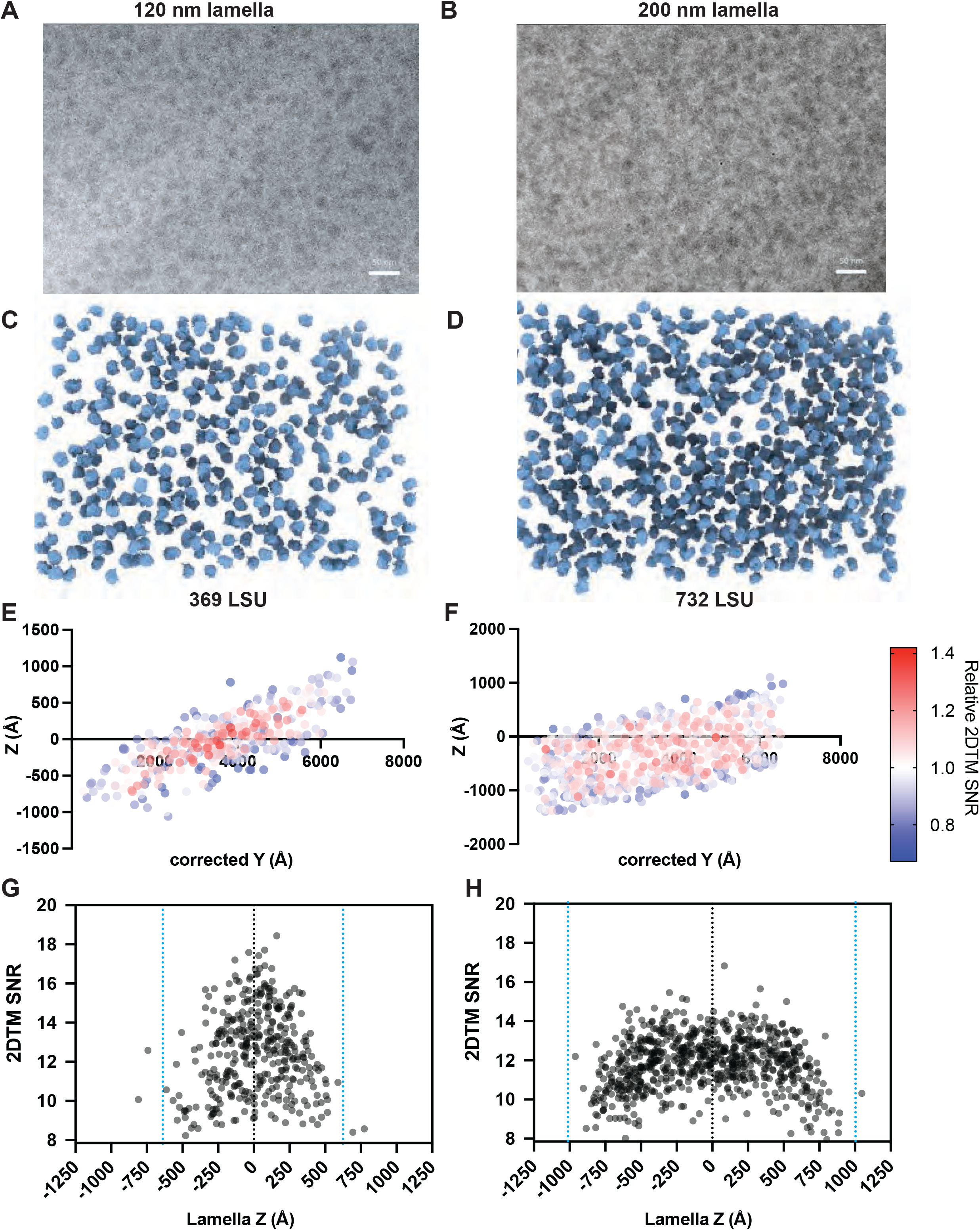
Visualization of yeast cytoplasmic ribosomes in 3D with 2DTM. **(*A*)** An electron micrograph of the yeast cytoplasm in a 120 nm region of a lamella. Scale bars in A) and B) represent 50 nm. **(*B*)** As in A), showing a 200 nm lamella. **(*C*)** Significant large ribosomal subunits (LSUs) located in 3D in the image in A) with 2DTM. **(*D*)** As in C), showing the results for the image in B). **(*E*)** Scatterplot showing a side view of the LSUs in A). The color coding indicates the 2DTM SNR of each significant detection relative to the mean 2DTM SNR. The z-coordinate represents the position of each target relative to the microscope defocus plane. **(*F*)** As in E), showing the results from the image in B). **(*G*)** Scatterplot showing the 2DTM SNR of each detected LSU in the image in A), as a function of z-coordinate relative to the lamella. **(*H*)** As in G), showing the z-coordinate relative to the center of the lamella of each LSU detected in the image shown in B).

We estimated the z-coordinate of each LSU relative to the image defocus plane with 2 nm precision (**Fig. 1*E* and *F***, see Methods). We found that the LSUs were located in a slab oriented at an angle of ∼6-11° relative to the defocus plane, consistent with the milling angle relative to the grid surface (**Fig. 1*C-F***). The 2DTM SNRs of LSUs were noticeably lower at the edge of the lamellae than at the center and did not correlate with defocus (**Fig. 1*E* and *F***), indicating that this is unlikely to be the result of defocus estimation error. We used the tilt axis and angle estimated from the contrast transfer function (CTF) fit (18, 19) to adjust the coordinate frame to reflect the position of each LSU relative to the lamella center (**Fig. 1*G* and *H***).

On average, the 2DTM SNRs were higher in the center and lower towards the surface in all lamellae examined (**Fig. 1 *G* and *H, SI Appendix*, Fig. S2**). The maximum 2DTM SNR decreased with increasing lamella thickness (***SI Appendix*, Fig. S1*B***) as observed previously (15–18). However, we observed a different 2DTM SNR profile as a function of z-coordinate in regions of different thicknesses. The 2DTM SNRs in ≤ ∼150 nm thick lamellae increased towards the center of the lamella (e.g.: **Fig. 1*G***), while in ≥ ∼150 nm thick lamellae they reached a plateau (e.g.: **Fig. 1*H***). This is consistent with decreased structural integrity of LSUs close to each lamella surface.

### Quantification of the damage profile reveals damage up to ∼60 nm from each lamella surface

To assess the depth of the damage we focused on images of 200 nm lamellae because we were able to detect targets throughout most of the volume, and both the number and 2DTM SNRs reached a plateau in the center, indicating that there is zone of minimal damage. In seven images of 200 nm lamellae, we calculated the mean 2DTM SNR in bins of 10 nm from the lamella surface and divided this by the undamaged SNR (*SNR*_*u*_), defined as the mean 2DTM SNR of the targets between 90 and 100 nm from the lamella surface. Both the relative 2DTM SNR (**Fig. 2*A***) and the number of LSUs detected (**Fig. 2*B***) increased as a function of distance from the lamella surface. The lower number of detected LSUs at the lamella surface is likely a consequence of targets having a 2DTM SNR that falls below the chosen 2DTM SNR threshold of 7.85 at which we expect a single false positive per image (15). Consistently, in each of the bins >60 nm from the lamella surface (**Fig. 2*C***), the distribution of 2DTM SNRs was Gaussian and not significantly different from the undamaged bin (t-test P>0.05, ***SI Appendix* Table S1**). However, for each of the bins ≤60 nm from the lamella surface, the distribution shifts significantly (t-test P<0.0001, ***SI Appendix* Table S1)** to the left, i.e., lower SNR values (**Fig. 2*C***). This indicates that the structural similarity between target and template decreases closer to the lamella surface. We interpret this as a loss of target integrity due to FIB-milling damage up to ∼60 nm from the lamella surface.

**Figure 2:**
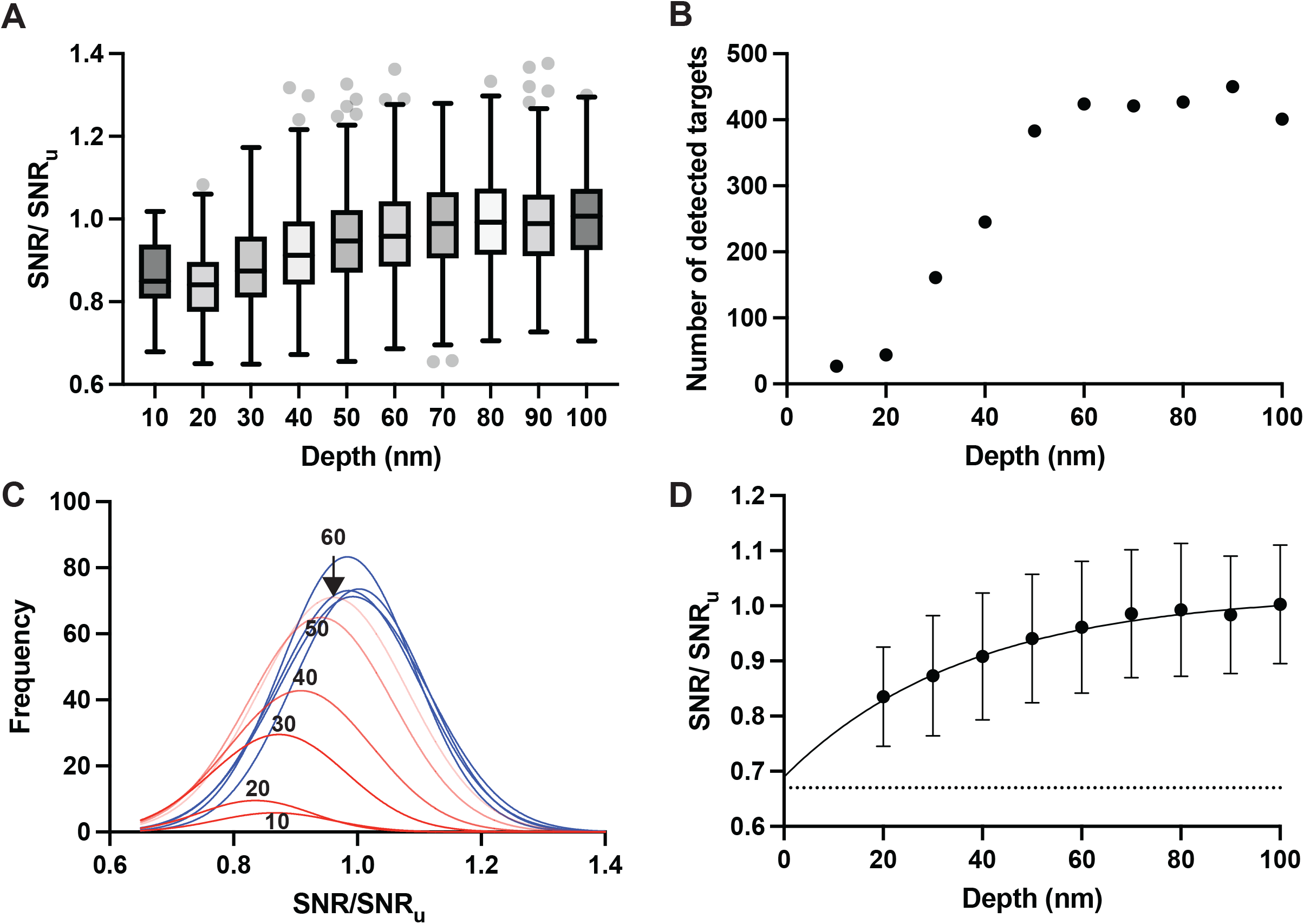
The number and 2DTM SNR values of detected LSUs increases as a function of distance from the lamella surface. **(*A*)** Boxplot showing the 2DTM SNR of LSUs at the indicated lamella depths, relative to the undamaged SNR (*SNR*_*u*_) in each image from 200 nm lamellae. Boxes represent the interquartile range (IQR), middle line indicates the median, whiskers represent 1.5x IQR and dots represent values outside of this range. **(*B*)** Scatterplot showing the number of detected targets in the indicated z-coordinate bins. **(*C*)** Gaussian fits to the distribution of 2DTM SNRs for LSUs identified in z-coordinate bins of 10 nm. Red indicates populations with means significantly different from the mean in the center of the lamella. Blue indicates that the mean in a bin is not significantly different from the mean in the center. Fitting statistics are indicated in ***SI Appendix*, Table S1. (*D*)** Scatterplot showing the mean change in 2DTM SNR relative to *SNR*_*u*_ at the indicated depths relative to the lamella surface estimated from the Gaussian fits in ***C***). The line shows the exponential fit (R^2^ = 0.99). Error bars indicated the standard deviation from the Gaussian fit.

We found that the change in the mean 2DTM SNR at a particular depth relative to the lamella surface (*d*) relative to *SNR*_*u*_ can be described by an exponential decay function:

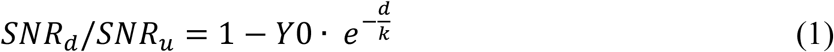

where *Y*0 is the relative damage at the lamella surface and *k* is the damage constant. A least-squares fit gave values of *Y*0 = 0.31 and *k* = 37.03 nm (R^2^=0.99) (**Fig. 2*D***). Since *SNR*_*d*_/*SNR*_*u*_ represents the remaining signal the exponential model indicates a steep decline in damage in the first ∼10-20 nm from the lamella surface, possibly explaining why few LSUs were detected in this range.

The observed damage profile was absent in images of unmilled *Mycoplasma pneumoniae* cells, confirming that the observed pattern results from FIB-milling and is not a result of error in the z-estimation in 2DTM (***SI Appendix*, Fig. S4 and 5**).

### Mechanism of FIB-milling damage

To characterize the mechanism of FIB-milling damage we compared its profile to the damage introduced by exposure to electrons during cryo-EM imaging. Cryo-EM imaging causes radiation damage, introducing differences between the template and the target structure that are more pronounced at high spatial frequencies (20). To measure radiation damage, we generated a series of low-pass filtered templates with a sharp cut-off at different spatial frequencies and calculated the change in the 2DTM SNR of each identified large ribosomal subunit as a function of electron exposure relative to exposure to 20 electrons/Å^2^ (**Fig. 3*A* and *B***). We find that the 2DTM SNR of templates low-pass filtered to between 1/10 and 1/7 Å^-1^ increases with increasing exposure. The 2DTM SNRs of templates low-pass filtered with a cut-off at higher resolutions begin to decrease with increasing exposure (**Fig. 3*A* and *B***). Templates filtered to 1/5 Å^-1^ have a maximum 2DTM SNR at 32 electrons/Å^2^ while templates filtered to 1/3 Å^-1^ have a maximum 2DTM SNR at 28 electrons/Å^2^ (**Fig. 3*B***).

**Figure 3:**
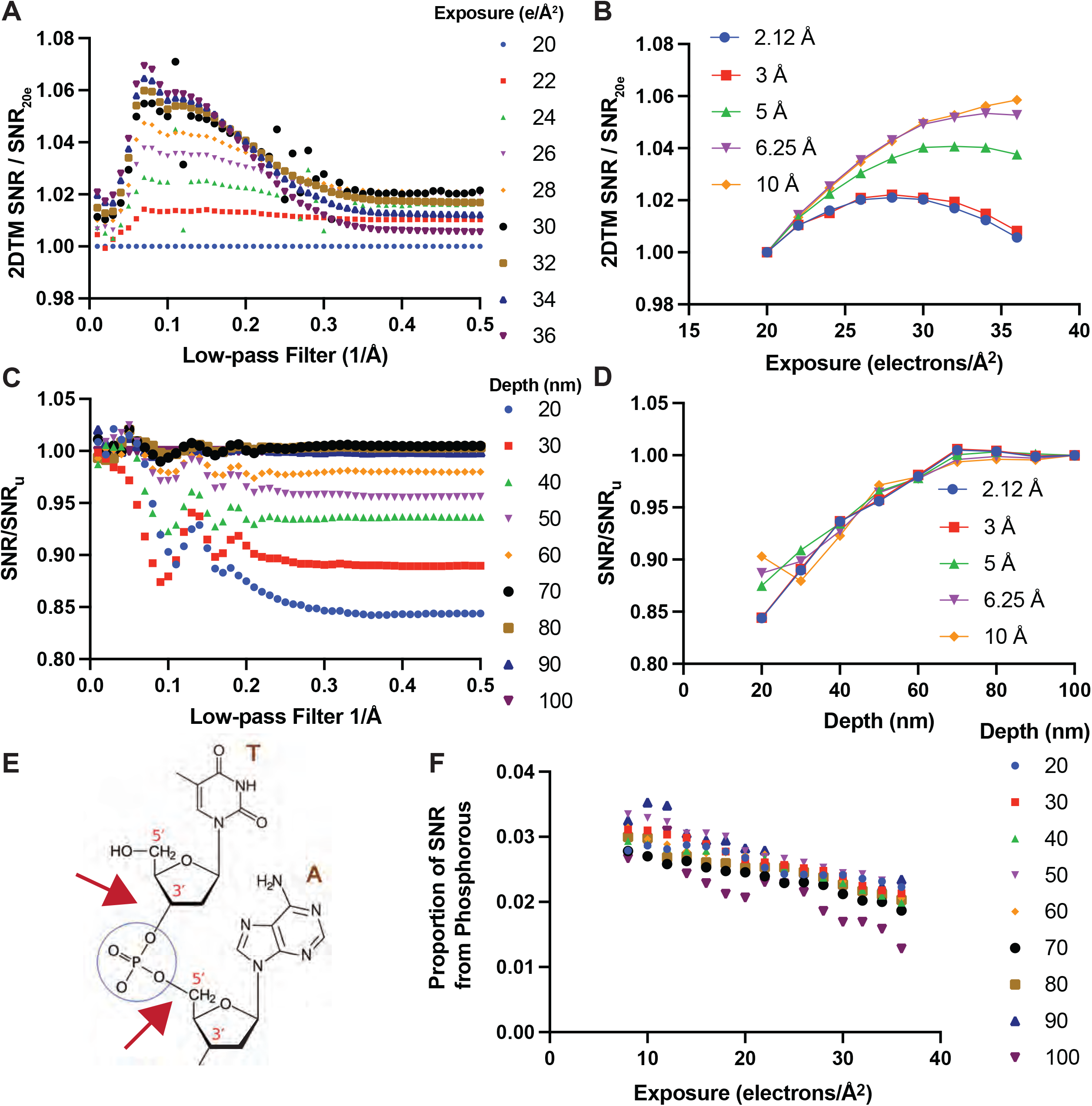
The mechanism of FIB-milling damage is distinct from radiation damage during cryo-EM imaging. **(*A*)** Plot showing the change in 2DTM SNR with the template low-pass filtered to the indicated spatial frequency in images collected with the indicated number of electrons/Å^2^ relative to the same filtered template with 20 electrons/Å^2^. **(*B*)** Plot showing the change in 2DTM SNR as a function of electron exposure of templates low-pass filtered to the indicated spatial frequency. **(*C*)** As in A), showing the change in the 2DTM SNR in the indicated lamella z-coordinate bins relative to the SNR in the undamaged bin (*SNR*_*u*_). **(*D*)** Plot showing the change in 2DTM SNR for templates low-pass filtered to the indicated spatial frequencies as a function of lamella z-coordinate bins. **(*E*)** Diagram showing a segment of an RNA strand of two nucleotides. The blue circle designates the phosphate; the two red arrows indicate the location of the backbone phosphodiester bonds. **(*F*)** Plot showing the relative contribution of template phosphorous atoms to the 2DTM SNR relative to the full-length template at the indicated exposure without dose weighting calculated using equation (6).

To estimate the extent of FIB-milling damage on different spatial frequencies we binned detected targets by lamella depth and calculated *SNR*_*d*_/*SNR*_*u*_. We found that for templates filtered to < 1/5 Å^-1^, *SNR*_*d*_/*SNR*_*u*_ fluctuated for targets detected further from the lamella center. This is likely due to differences in the defocus position that result in some of the targets having weak contrast (CTF close to zero) and therefore not contributing meaningful signal at different spatial frequencies relative to targets in the center of the lamella. For templates filtered to > 1/5 Å^-1^ the profile was similar between the different bins and approximately constant across spatial frequencies (**Fig. 3*C* and *D***). This is consistent with a model in which the FIB-damaged targets have effectively lost a fraction of their structure, compared to undamaged targets, possibly due to displacement of a subset of atoms by colliding ions.

Radiation damage of nucleic acids has been well documented with one of the most labile bonds being the phosphodiester bond in the nucleic acid backbone (21) (**Fig. 3*E***). We observed an accelerated loss of signal from phosphorous atoms relative to the average loss of signal for the whole template as a function of electron exposure (**Fig. 3*F***). This is consistent with the phosphorous atoms being more mobile due to breakage of phosphodiester linkages in response to electron exposure. We did not observe a consistent difference in the accelerated loss of signal from phosphorous in the lamella z-coordinate groups (**Fig. 3*F***). This indicates that the mechanism for FIB-milling damage is distinct from the radiation damage observed during cryo-EM imaging.

### Sample thickness limits 2DTM SNR more than FIB-milling damage

Above we report that using the most common protocol for cryo-lamella generation by LMIS Ga^+^ FIB-milling introduces a variable layer of damage up to 60 nm from each lamella surface. Lamellae for cryo-EM and cryo-ET are typically milled to 100-300 nm, meaning that the damaged layer comprises 50-100% of the volume. Thicker lamellae will have a lower proportion of damaged particles. However, thicker lamellae will also suffer from signal loss due to the increased loss of electrons due to inelastic scattering and scattering outside the aperture, as well as the increased number of other molecules in the sample contributing to the background in the images. For a target inside a cell, the loss of 2DTM SNR with increasing thickness has been estimated as (16):

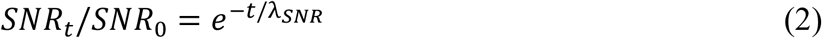

where *t* denotes the sample thickness, *SNR*_0_ is the 2DTM SNR in the limit of an infinitesimally thin sample, and the decay constant λ_*SNR*_ = 426 nm. Optimal milling conditions for high resolution imaging of FIB-milled lamellae will therefore need to strike a balance between lamella thickness and FIB damage.

To assess the relative impact of these two factors on target detection with 2DTM we plotted the proportional loss in signal due to electrons lost in the image, and background (**Fig. 4**, red curve). We can estimate the average loss of 2DTM SNR, 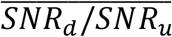, due to FIB-milling damage from the product of the loss (Eq. (1)) from both surfaces:

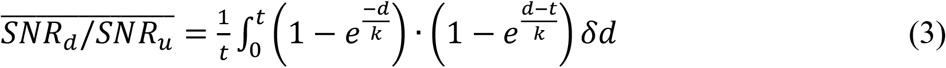

**Figure 4:**
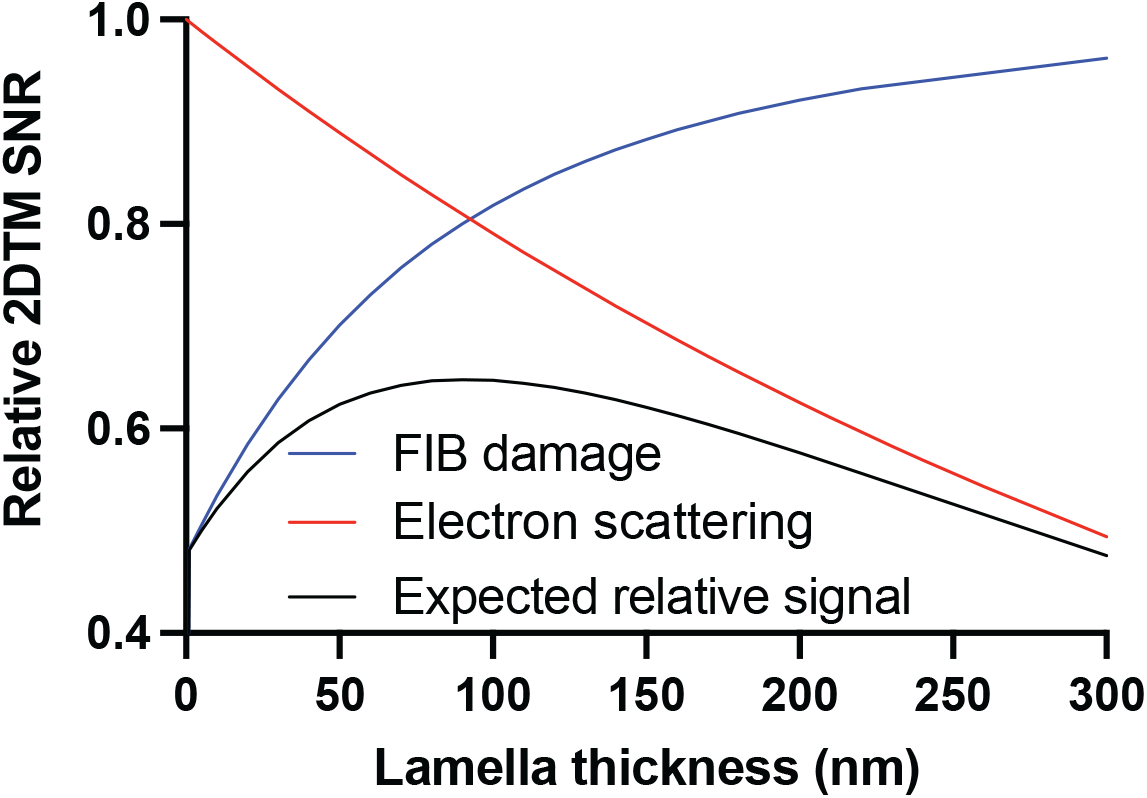
Signal loss due to increased electron scattering in thicker samples outweighs the effect of FIB damage on LSU 2DTM SNRs. Plot showing the expected signal recovery in lamellae of indicated thickness as a function of signal loss due to electron scattering (red curve), FIB-damage (blue curve) and their product (black curve).

Combining these two sources of signal loss gives the expected overall 2DTM SNR as a function of sample thickness (**Fig. 4**, black curve):

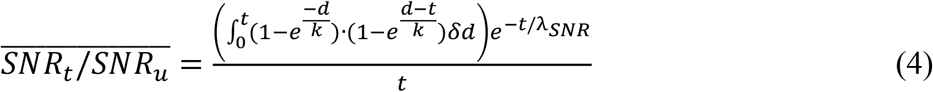

This model predicts that in samples thicker than ∼90 nm, the relative loss in the signal due to the loss of electrons contributing to the image, as well as molecular overlap is greater than the relative change due to FIB-milling damage (**Fig. 4*A***). In lamellae thinner than 90 nm, however, FIB-milling damage will dominate and negate any benefit from further thinning. The difference between the expected signal loss given by Eq. (4) and signal loss solely from lost electrons and molecular overlap represents the potential gain if FIB-milling damage could be avoided. Without FIB damage, the potential improvement in 2DTM SNR would be between ∼10% in 200 nm lamellae and ∼20% in 100 nm lamellae (**Fig. 4**). The model in Eq. (4) ignores the variable degree of damage expected to occur across LSUs that we used as probes to measure damage, and that have a radius of ∼15 nm. However, the resulting error in the measured damage constant *k* (Eq. (1)) is expected to be small since *k* (∼37 nm) significantly exceeds the LSU radius and hence, the variable damage can be approximated by an average damage uniformly distributed across the target.

We also expect that the number of detected targets will be reduced by FIB-milling damage. The number of detected LSUs was variable across lamellae, likely due to biological differences in local ribosome concentration. In undamaged parts of a subset of 200 nm thick lamellae, we identified ∼425 LSU in z-coordinate intervals of 10 nm. If this density were maintained throughout the lamella, we would expect to detect ∼40% more targets in these lamellae.

We conclude that FIB-damage reduces the number and integrity of detected targets but that signal loss due to electrons lost to the image, as well as background from overlapping molecules is a greater limiting factor for target detection and characterization with 2DTM than FIB-milling damage in lamellae thicker than ∼90 nm. These data agree with other empirical observations that thinner lamellae are optimal for recovery of structural information and generation of high-resolution reconstructions.

It may be possible to restore signal in images otherwise lost to inelastic scattering using Cc-correctors (22). This would be particularly impactful for thick samples such as FIB-milled cellular lamellae. With the use of a Cc-corrector, the signal loss in thick samples would be reduced and FIB-milling damage may become the main limiting factor for *in situ* structural biology.

## Discussion

Ga^+^ LMIS FIB-milling is currently the preferred method for generating thin, electron-transparent cell sections for *in situ* cryo-EM. We use 2DTM to evaluate the structural integrity of macromolecules in FIB-milled lamellae and provide evidence that FIB-milled lamellae have a region of structural damage to a depth of up to 60 nm from the lamella surface. By evaluating the relative similarity of a target molecule to a template model, 2DTM provides a sensitive, highly position-specific, single particle evaluation of sample integrity.

### 2DTM SNRs provide a readout of sample integrity and image quality

Changes to the 2DTM SNR provide a read-out of the relative similarity of a target molecule to a given template. We have previously shown that relative 2DTM SNRs discriminate between molecular states and can reveal target identity (17, 18). In this study we show that changes in 2DTM SNRs can also reflect damage introduced during FIB-milling and radiation damage introduced during cryo-EM imaging. Previous attempts to measure FIB-damage have relied on visual changes in the sample near the surface. These changes are difficult to quantify in terms of damage, and they could in part be caused by other mechanisms such as ice accumulation after milling (23). Argon plasma FIB-damage has been assessed by estimating B-factors of subtomogram averages, comparing averages of particles at less than and greater than 30 nm from the lamella surface, which lacks positional precision (24). The 2DTM SNR represents a more quantitative metric to assess sample integrity.

2DTM SNRs have also been used as a metric to assess image quality (25) and the fidelity of simulations (26). 2DTM therefore, represents a sensitive, quantitative, and versatile method to measure the dependence of data quality on sample preparation and data collection strategies, as well as new hardware technologies and image processing pipelines. Tool and method developers could use standard datasets and 2DTM to rapidly and quantitatively assess how any changes to a pipeline affect data quality.

### Estimating errors in z-coordinates and thickness

The z-coordinates of each LSU were determined by modulating the template with a CTF corresponding to a range of defoci and identifying the defocus at which the cross-correlation with the 2D projection image was maximized (15). This quantification relies on an accurate estimate of defocus. The error in the z-coordinates determined this way was estimated to be about 60 Å (17). However, it is unlikely that these errors explain the observed decrease in 2DTM SNRs of LSUs near the edge of the lamellae because (1) the reduction in 2DTM SNRs correlates strongly with the z-coordinate within the lamella, and (2), we did not observe a consistent decrease in the number of detected LSUs (*SI Appendix*, **Fig. S4*A***), or their 2DTM SNRs (*SI Appendix*, **Fig. S4*B***) as a function of z-coordinate in images of unmilled *M. pneumoniae* cells (17).

Undulations at the lamella surface caused by curtaining or other milling artifacts could contribute to the reduced number of ribosomes detected near the lamella surface. We aimed to minimize the effect of curtaining in our analysis by calculating the lamella thickness in 120 × 120 pixel (127.2 × 127.2 Å) patches across an image and limiting our analysis to images with a thickness standard deviation of less than 20 nm. The curtaining on the remaining lamellae cannot account for the reduced particle integrity towards the lamella surface.

### Possible mechanisms of FIB-milling damage

We find evidence for FIB-milling damage consistent with an exponential decay of the amount of damage as a function of distance from the lamella surface, as measured by the 2DTM SNR. Unlike electron radiation damage, FIB-damage 1) causes a reduction in the total signal and does not preferentially affect higher spatial frequencies contributing to the 2DTM SNR calculation, and 2), unlike electron beam radiation damage, it does not preferentially affect the phosphodiester bond in the RNA backbone. This suggests that different mechanisms are responsible for the damage caused by high-energy electrons and ions.

At the energy ranges used for FIB-milling, the interactions between the bombarding ion and sample atoms can be modelled as a cascade of atom displacements resulting from the transfer of momentum from the incident Ga^+^ ions to the sample atoms (13). Atoms involved in the cascade will be displaced while the position of other atoms will not change. This is consistent with our observation that FIB damage decreases the LSU target signal overall without changing the relative contribution from different spatial frequencies. Further study is required to test this hypothesis and investigate the mechanism of FIB-milling damage in more detail.

SRIM simulations predict implantation of Ga^+^ up to ∼25 nm into the sample (5, 12). This implies that the damage deeper in the sample is caused by secondary effects, possibly reflecting displaced sample atoms that were part of the collision cascade. Interestingly, we observe a different pattern of particles within 20 nm of the lamella surface (**Fig. 3*C* and *D***). One possible explanation is that implanted Ga^+^ ions cause additional damage. However, SRIM simulations have been shown previously not to account for the full intensity profile of a Ga^+^ beam, and poorly match with experiment especially at low beam currents (27). Moreover, the use of a protective organo-platinum layer during FIB-milling, as done in our experiments, will further change the effective profile of the beam acting on the sample (28). Further work is required to connect the quantification of particle integrity with the implantation of Ga^+^ ions during biological lamella preparation.

### Implications for generating high resolution reconstructions from FIB-milled samples

We have shown that particles on the edge of a lamella have reduced structural integrity relative to particles near the center of the lamella (**Fig. 1 and 2**). We found that FIB-milling damage reduces the total 2DTM SNR and that >20 nm from the lamella surface the rate of signal loss is similar at different spatial frequencies, in contrast to radiation damage during cryo-EM imaging (**Fig. 3**). The practical implication of this finding is that particles >30 nm from the lamella surface can be included during subtomogram averaging without negatively affecting the resolution of the reconstruction, provided they can be accurately aligned. We also predict that more particles will be required relative to unmilled samples. This is consistent with the observation that more particles <30 nm from the lamella surface are required to achieve the same resolution relative to >30 nm from the lamella surface from Argon plasma FIB-milled lamellae (24).

Due to the small number of particles detected within 10 nm of the lamella surface, these particles were not examined in more detail. Since ribosomes are ∼25 nm in diameter, it is likely that these particles are more severely damaged compared to particles further away from the surface. 2DTM relies on high-resolution signal and therefore excludes more severely damaged particles that may be included using a low-resolution template matching approach, such as 3D template matching used typically to identify particles for subtomogram averaging. We therefore advise excluding particles detected within 10 nm of the lamella surface.

### Alternate methods for the preparation of thin cell sections

FIB damage reduces both the number of detected targets and the available signal per target. However, the damaged volume still contributes to the sample thickness, reducing the usable signal by 10-20% in lamellae of typical thicknesses (**Fig. 4**). Therefore, it would be advantageous to explore other strategies for cell thinning.

Plasma FIBs allow different ions to be used for milling, and this may change the damage profile (29). Larger atoms such as Xenon will have a higher sputtering yield and may result in reduced lamella damage, as has been demonstrated for milling of silicon samples (30, 31). The 2DTM-based approach described here provides a straightforward way to quantify the relative damaging effects of different ion species by generating curves as shown in Fig. 4.

CEMOVIS generates thin sections using a diamond knife rather than high energy ions and would therefore not introduce radiation damage (2). It is unclear how the large-scale compression artifacts introduced by this method affect particle integrity (3). CEMOVIS has the additional benefit of being able to generate multiple sections per cell and thereby enable serial imaging of larger cell volumes. If the compression artifacts are unevenly distributed throughout a section, leaving some regions undistorted, automation could make CEMOVIS a viable strategy for structural cell biology in the future.

To retain the benefits of fast, reliable, high-throughput lamella generation with cryo-FIB milling, strategies to remove the damaged layer should be explored. Possibilities include polishing the final ∼50 nm from each lamella surface with a low energy (∼5 kV) beam, which has the advantage of being easily implementable using the current configuration of most cryo-FIB-SEMs.

## Materials and Methods

### Yeast culture and Freezing

*Saccharomyces cerevisiae* strains BY4741 (ATCC) colonies were inoculated in 20 mL of YPD, diluted 1/5 and grown overnight at 30 °C with shaking to mid log phase. The cells were then diluted to 10,000 cells/mL, treated with 10 µg/mL cycloheximide (Sigma) for 10 mins at 30 °C with shaking and 3 µL applied to a 2/1 or 2/2 Quantifoil 200 mesh SiO_2_ Cu grid, allowed to rest for 15 s, back-side blotted for 8 s at 27 °C, 95% humidity followed by plunge freezing in liquid ethane at –184 °C using a Leica EM GP2 plunger. Frozen grids were stored in liquid nitrogen until FIB-milled.

### FIB-milling

Grids were transferred to an Aquilos 2 cryo-FIB/SEM, sputter coated with metallic Pt for 10 s then coated with organo-Pt for 30 s and milled in a series of sequential milling steps using a 30kV Ga+ LMIS beam using the following protocol: rough milling 1: 0.1 nA rough milling 2: 50 pA lamella polishing: 10 pA at a stage tilt of 15° (milling angle of 8°) or 18° (milling angle of 11°).

### Cryo-EM data collection and image processing

Cryo-EM data was collected following the protocol described in (18) using a Thermo Fisher Krios 300 kV electron microscope equipped with a Gatan K3 direct detector and Gatan energy filter with a slit width of 20 eV at a nominal magnification of 81,000x (pixel size of 1.06 Å^2^) and a 100 *μ*m objective aperture. Movies were collected at an exposure rate of 1 e^-^/Å^2^/frame to a total dose of 50 e^-^/Å^2^ (Dataset 1) or 30 e^-^/Å^2^ (Dataset 2) with correlated double sampling using the microscope control software SerialEM (32).

Images were processed as described previously (18). Briefly, movie frames were aligned using the program *unblur* (33) in the *cis*TEM GUI (34) with or without dose weighting using the default parameters where indicated in the text. Defocus, astigmatism, and sample tilt were estimated using a modified version of CTFFIND4 (17, 19) in the *cis*TEM GUI (34). Images of cytoplasm were identified visually for further analysis. Images visually containing organelles were excluded. Images of 3D densities and 2DTM results were prepared in ChimeraX (35).

### 2DTM

The atomic coordinates corresponding to the yeast LSU from PDB: 6Q8Y (36) were used to generate a 3D volume using the *cis*TEM program *simulate* (26) and custom scripts as in (21). 2DTM was performed using the program *match_template* (20) in the *cis*TEM GUI (34) using an in-plane search step of 1.5° and an out-of-plane search step of 2.5°. Significant targets were defined as described in (15) and based on the significance criterion described in (16). The coordinates were refined using the program *refine_template* (18) in rotational steps of 0.1° and a defocus range of 200 Å with a 20 Å step (2 nm z-precision). The template volume was placed in the identified locations and orientations using the program *make_template_result* (17) and visualized with ChimeraX (35).

To generate the results in Fig. 3A-D, we applied a series of sharp low-pass filters in steps of 0.01 Å^-1^ to the template using the e2proc3d.py function in EMAN2 (37). We used the locations and orientations from the refined 2DTM search with the full-length template to recalculate the 2DTM SNR with each modified template using the program *refine_template* (18) by keeping the positions and orientations fixed. The normalized cross-correlation was determined by dividing the SNR calculated with each low-pass filtered template to the SNR of the full-length template for each target.

### Calculation of tilt and coordinate transform

We used python scripts to extract the rotation angle and tilt from the *cis*TEM (34) database generated using the tilt-enabled version of the program CTFFIND4 (18, 19), perform a coordinate transform to convert the 2DTM coordinates to the lamella coordinate frame and plot the 2DTM SNR as a function of lamella z-coordinate.

### Calculation of sample thickness and depth

We estimated the lamella thickness per image by first summing the movie frames without dose weighting using the EMAN2 program, *alignframe*s (37), then calculating the average intensity of a sliding box of 100 × 100 pixels (*I*) relative to the same area of an image collected over vacuum (*I*_*o*_). We then used the mean free path for electron scattering (*λ*) of 283 nm (16) to estimate the local sample thickness *t*_*i*_ using the Beer-Lambert law (38):

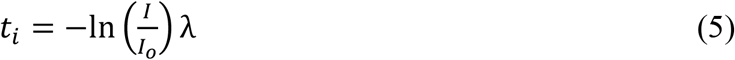

The sample thickness was determined by taking the mean across the image. Only images with a standard deviation of <20 nm across the image were included for estimation of the damage profile (**Fig. 2*B***). The depth of each LSU relative to the lamella surface was calculated by assuming that the LSUs are evenly distributed in z and defining the median lamella z coordinate as the lamella center (e.g.: **Fig. 1*G,H* and S2**).

### Measuring change in signal with electron exposure

We compared the change in the 2DTM SNR of each individual LSU as a function of electron exposure at different positions relative to the edge of the lamella in bins of 10 nm. We used the locations and orientations of LSUs identified in dose-filtered images exposed to 50 e^-^/Å^2^ to assess the correlation at the same locations and orientations in different numbers of unweighted frames corresponding to total exposures of 8-36 e^-^/Å^2^.

To calculate the relative contribution of phosphorous to the 2DTM SNR all phosphorous atoms in the PDB file were deleted and a template was generated as described above without re-centering so that it aligned with the full-length template. We used the locations and orientations from the refined 2DTM search with the full-length template for each exposure to calculate the 2DTM SNR with the template lacking phosphorous (*SNR*_Δ*P*_) using the program *refine_template* (18) and keeping the positions and orientations fixed. The relative contribution of phosphorous atoms to the 2DTM SNR (*SNR*_*P*_) at each exposure was calculated using the following equation:

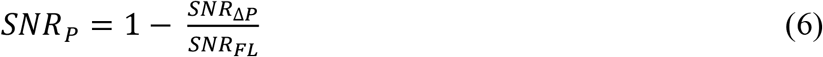

## Acknowledgements

The authors thank Johannes Elferich, Ximena Zottig and other members of the Grigorieff lab (UMass Chan), Russo lab (MRC LMB) and de Marco lab (Monash) for helpful discussions. We are also grateful for the use of and support from the cryo-EM facilities at Janelia Research Campus and UMass Chan Medical School.

## Figure Legends

**Figure S1:**
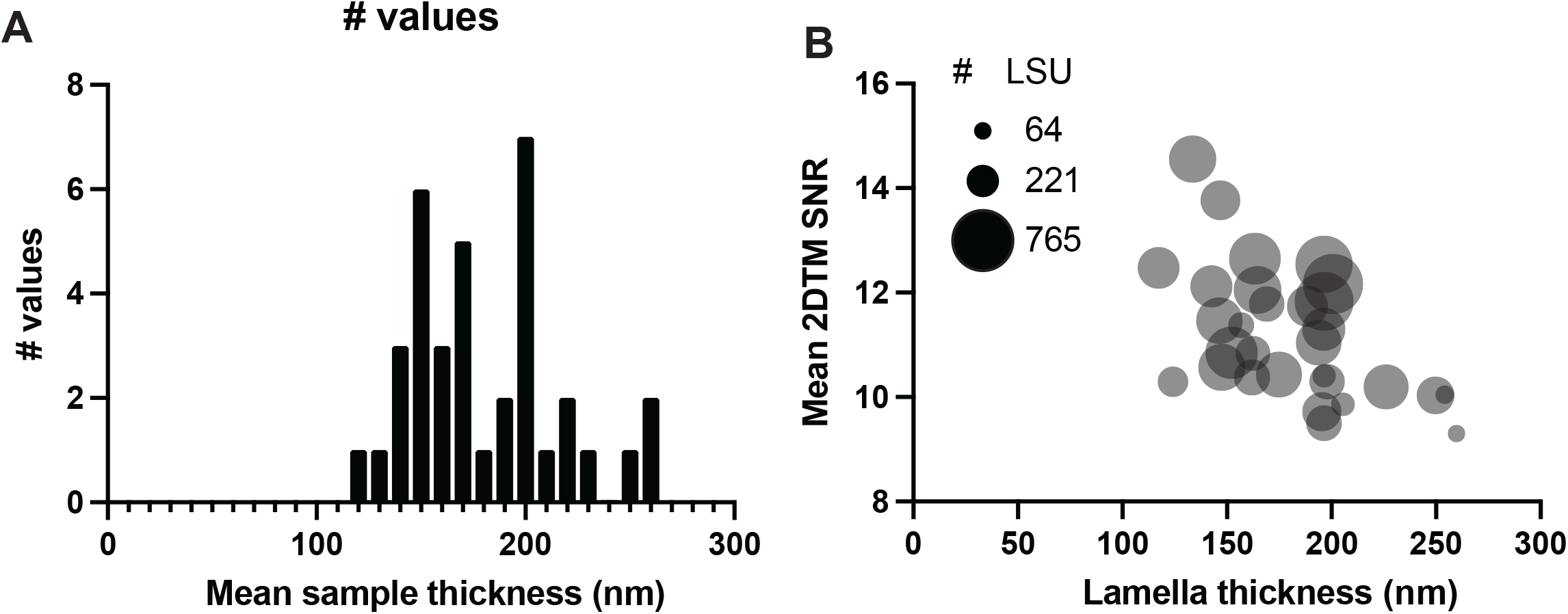
**(*A*)** Histogram showing the distribution of lamella thickness calculated using the Beer-Lambert law. **(*B*)** Scatterplot showing the mean 2DTM SNR of the detected LSUs in each image as a function of local lamella thickness. The point size indicates the number of LSUs detected in the image.

**Figure S2:**
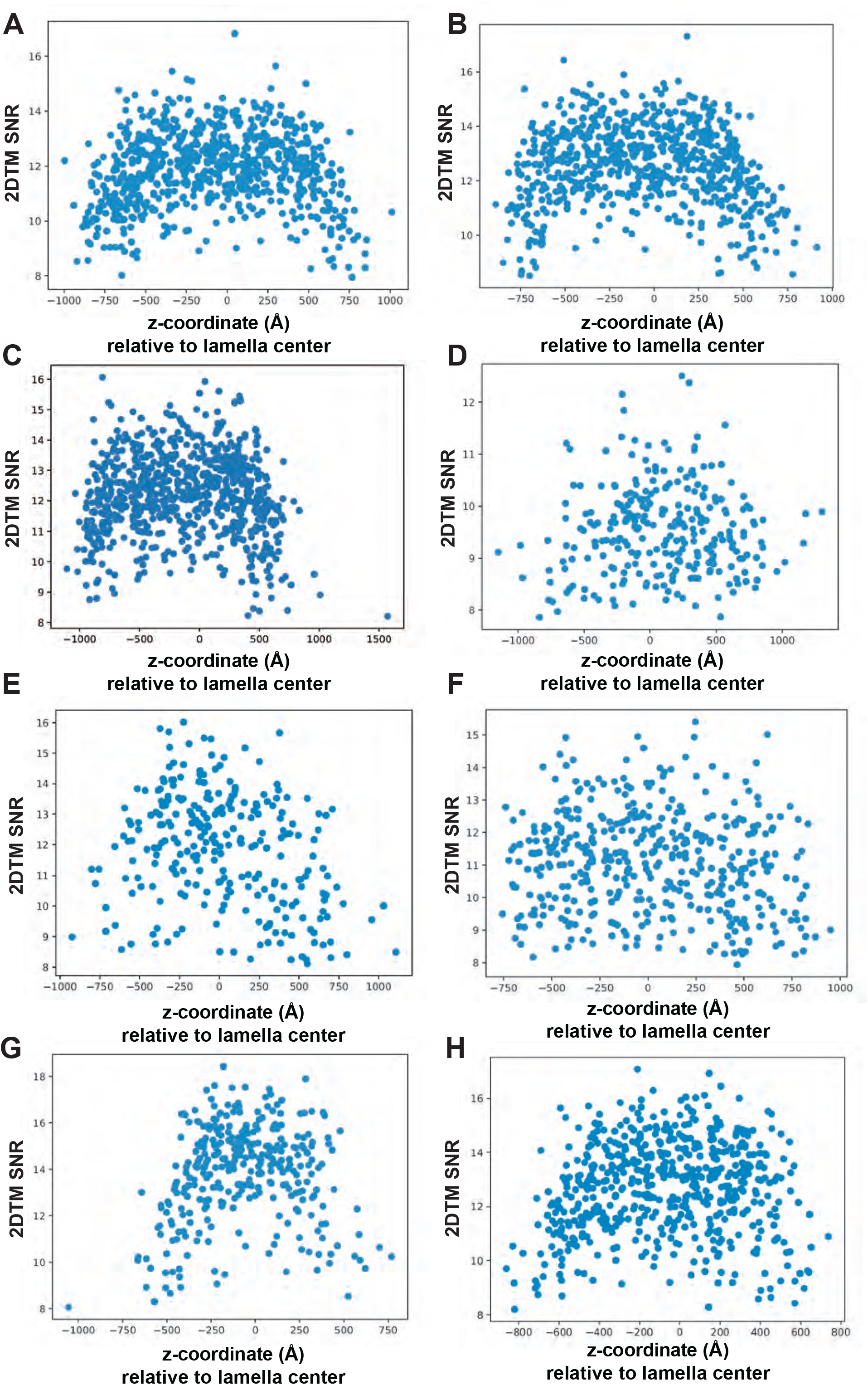
Representative plots showing the relationship between lamella z-coordinate and 2DTM SNR of LSU-detected targets in FIB-milled yeast lamellae.

**Figure S3:**
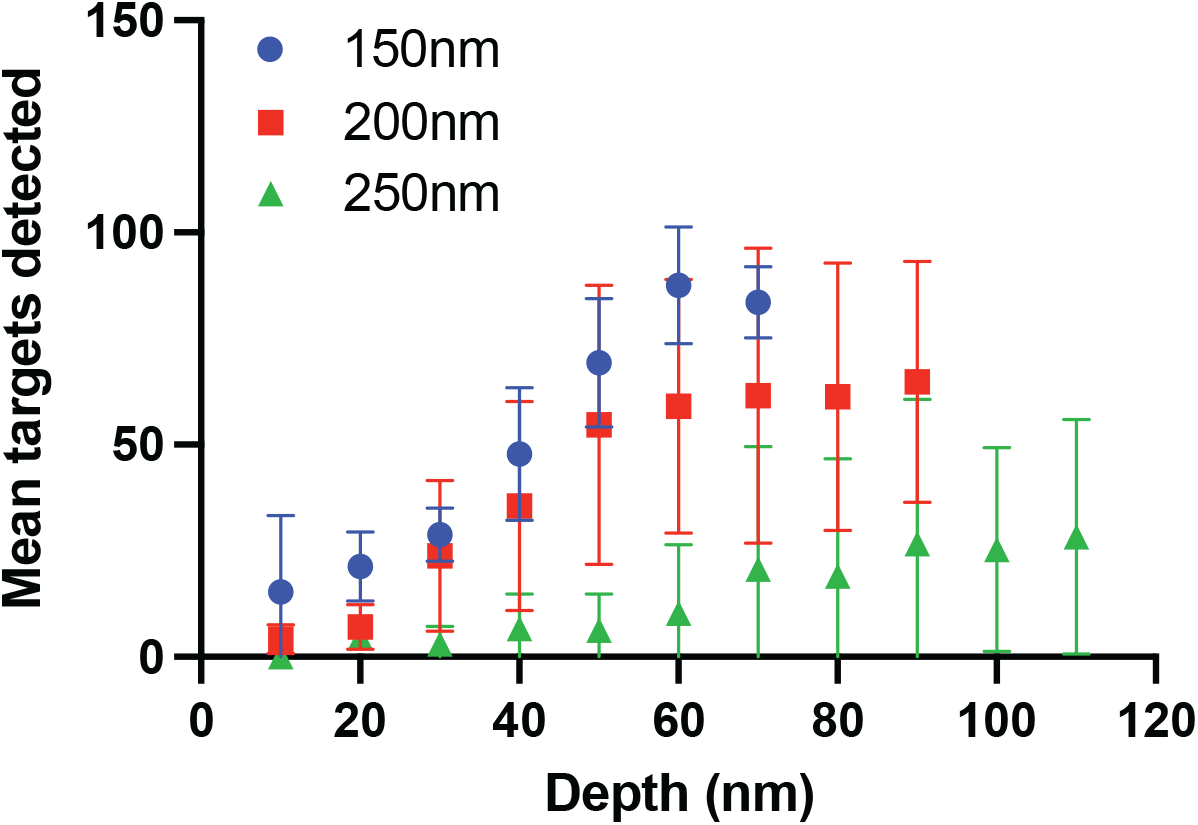
Scatterplot showing the mean number of targets detected in each lamella z-coordinate bin for lamellae of the indicated thickness. Error bars indicate the standard deviation

**Figure S4:**
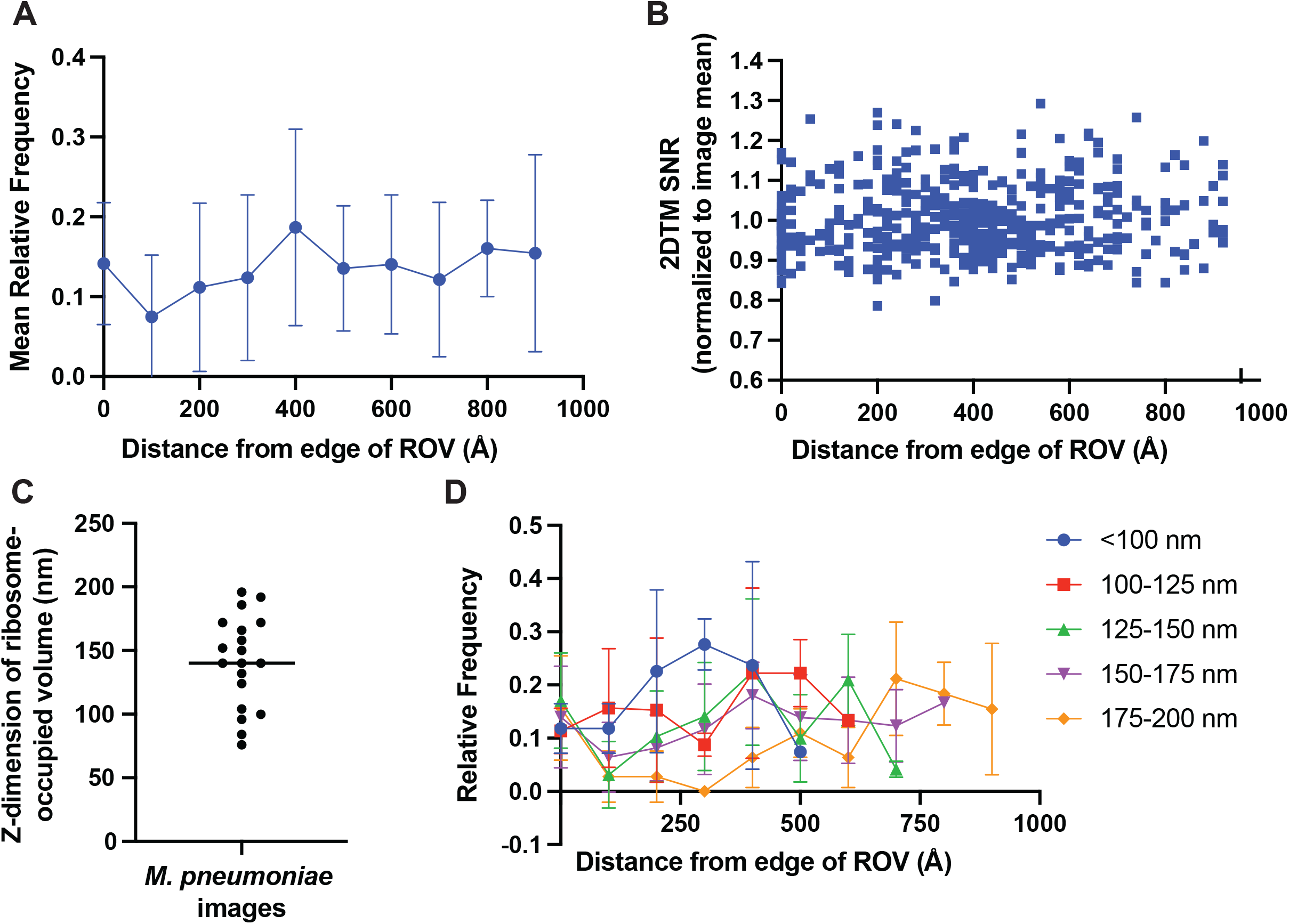
No consistent relationship between 2DTM SNRs and depth in unmilled *Mycoplasma pneumoniae* cells. **(*A*)** Scatterplot showing the mean proportion of LSUs identified in each bin as a function of distance from the edge of the ribosome occupied volume (ROV). **(*B*)** Scatterplot showing the 2DTM SNR relative to the image mean for each identified LSU as a function of distance from the edge of the ROV. **(*C*)** Plot showing the z-dimension of the ROV in images of *Mycoplasma pneumoniae* cells. **(*D*)** As in A), showing the results grouped by thickness.

**Figure S5:**
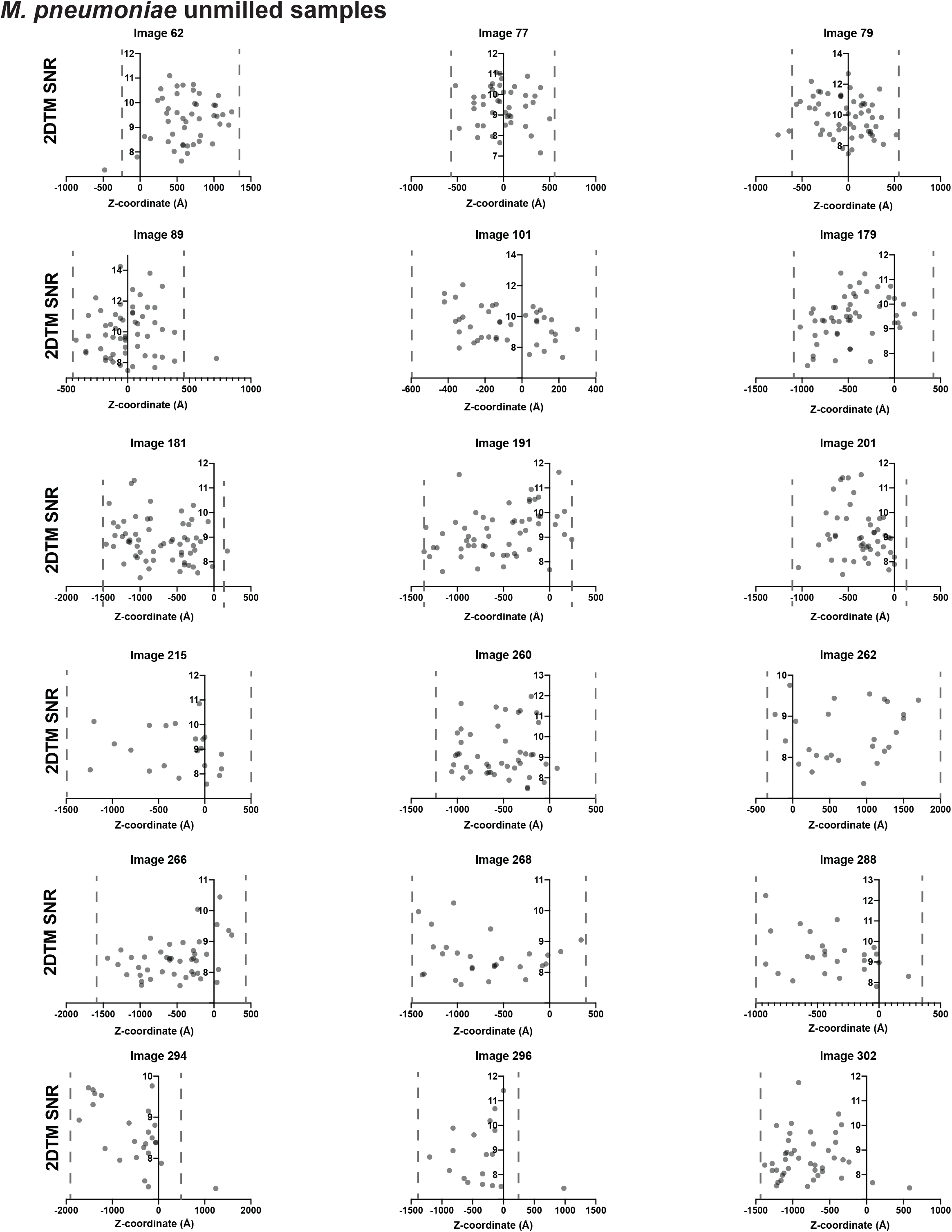
Scatterplots showing the 2DTM SNRs of LSU-detected targets in images of unmilled *Mycoplasma pneumoniae* cells as a function of z-coordinate relative to the defocus plane at x = 0.

---

**Table S1:** Gaussian fits to z-coordinate bins in Figure 2.

| Depth (nm) | Mean | SD | R squared | N | t-test vs 100 nm |
| --- | --- | --- | --- | --- | --- |
| **10** | 0.8664 | 0.09367 | 0.8148 | 27 | P<0.0001 |
| **20** | 0.8353 | 0.09015 | 0.9243 | 44 | P<0.0001 |
| **30** | 0.8733 | 0.1089 | 0.9591 | 161 | P<0.0001 |
| **40** | 0.9084 | 0.115 | 0.9345 | 245 | P<0.0001 |
| **50** | 0.9405 | 0.1167 | 0.989 | 383 | P<0.0001 |
| **60** | 0.9612 | 0.1193 | 0.9839 | 424 | P<0.0001 |
| **70** | 0.9859 | 0.1163 | 0.9644 | 421 | ns |
| **80** | 0.9926 | 0.1207 | 0.9703 | 427 | ns |
| **90** | 0.9837 | 0.1065 | 0.9676 | 450 | ns |
| **100** | 1.003 | 0.1079 | 0.9865 | 401 | _ |

